# Investigating the Impact of Measurement Variance on Gene Circuit Model Parameterization

**DOI:** 10.1101/2025.04.02.646175

**Authors:** Thales R. Spartalis, Wan Tang, Xun Tang

## Abstract

Ordinary differential equation (ODE)-based modeling is a powerful tool in the design and characterization of synthetic gene circuits. Despite its popularity, identifying the model parameters based off experimental measurement is a nontrivial task. In this study, we leverage cell-free experimental measurement of two RNA-based regulators to investigate the impact and the incorporation of measurement variance in the pair-wise squared error objective function used for ODE-model parameterization. Our findings suggest that while unweighted objective function and weighting by the inverse variance can provide reasonably accurate parameter estimation, weighing the objective function with the inverse stabilized variance could further improve the parameterization, by also capturing the system variance with a mitigated prediction variance.

## 2. Introduction

Mathematical modeling is a versatile tool in synthetic biology, from deciphering the complex metabolic pathways to the design of novel synthetic regulatory networks, and to the engineering of novel diagnostical and therapeutical solutions. [1–4] Synthetic gene circuits are composed by biomolecular components such as DNAs, RNAs, and proteins, and are engineered to exhibit specific functionalities by designing the interaction pathways among their constituent components, and with the environment. It is pivotal to understand the properties of a newly devised gene circuit to ensure the desired functionality before its practical implementation. Advances in experimental testbed such as the cell-free transcription-translation system, have significantly reduced the cost of experiment-based characterization of synthetic gene circuits with a controlled environment, while providing a means of high throughput experimental measurements. However, conducting extreme or customized-condition evaluation of the circuit property in experiments still presents a challenge, yet desirable. Coupled with cell-free experiments, mathematical modeling can facilitate the characterization of the circuit properties. By leveraging experimental gene expression data, scientists can infer interactions, reconstruct network edges, and describe system dynamics in complex biological systems, through mathematical expressions. This model-based approach complements experimental efforts to enable efficient quantification of system behavior, and cost-effective exploration of the underlying mechanisms that drive cellular functions. [1,5]

The dynamics of a synthetic gene circuit can be modeled with parametric models such as partial or ordinary differential equations, [6–12] and non-parametric models such as Markov state models, [13] artificial neural networks, [14] as deterministic or stochastic modeling. Arguably, deterministic ordinary differential equation (ODE)-based models are one of the simplest and most widely used model types for simulating gene circuit dynamics. ODE-based models are typically developed with fundamental understanding of the chemical reactions in the system, and describe the concentration evolution of the species of interest in the circuit with kinetic parameters, at the averaged cell-population level. While the accuracy of the ODE-based models heavily depends on model complexity, [15,16] and the correctness of knowledge on the reactions in the system, [17,18] the specific values of the kinetic parameters such as transcription, translation, degradation, and binding rate of the biomolecules in the system (e.g. ribosome and RNA polymerase binding rates), also play a decisive role in the model accuracy. [19] However, determining the unknown model parameters, a.k.a. model parameterization is a long-standing challenge, due to the model structure and properties (e.g. singularity and stiffness), parameter dimensionality and uncertainty, as well as limited experimental measurements. [20–23]

Parameterization of a gene circuit model is typically achieved by minimizing a cost function, which quantifies the discrepancy between the simulated and the experimental measurements. While it can be formulated in various forms, [20,24] summation of the pair-wise mean squared error (MSE) is one of the most widely used objective functions, given its simplicity and ease of understanding. Inverse variance weighted MSE is also proposed to account for measurement accuracy. However, findings suggest that the specific formulation of the MSE-based cost function, especially the weighting of the pair-wise error at different time points could have a significant impact on the accuracy of the estimated parameters. [25,26] In Ref. [25], the authors provided a detailed review on recent studies on the design of the error measure models (i.e. objective functions) and their corresponding performance.

While weighted cost function has attracted wide attentions in various fields, [27–29] the design of objective function, especially on simple weighted objective functions for synthetic gene circuits model parameterization is still under-investigated. One of the reasons is that the measurement variance could differ significantly from time to time and from system to system. [26] Here, to fill the knowledge gap and to explore the design rules for simple yet reliable parameter estimation for synthetic gene circuits modeling, we focus on two RNA-based regulators. Specifically, we focus on understanding the impact of experimental measurement variance on model parameterization, and how to incorporate such variance in the formulation of the objective function for improved model parameter identification. The study is organized as the following: 1) we first focus on a STAR system, and generate a simulated dataset to explore the objective function design guidelines, and 2) we then apply the developed guidelines to a CRISPRi system for validation, and 3) we conclude with discussion on the key findings, open questions and future work.

## 3. Methods

### STAR System and Synthetic Data Generation

We first focus on a small transcription activating RNAs (STAR) system, depicted in Fig. 1, same as in Ref. [30]. The STAR system turns on the originally transcription-off gene by opening the hairpin of the target gene to resume transcription. To simulate a simplified representation of this process, we adopted the same ODE model as in Ref. [30], which describes the following mechanism: 1) the STAR RNA is constitutively transcribed from the STAR DNA plasmid (*P*_*s*_), at a rate of *α*_*s*_; 2) instead of binding to the unfolded target mRNA strand, to simplify the model, the STAR RNA is modeled to directly bind the Target DNA plasmid (*P*_*y*_) to enable transcription of the target gene, at a rate *β*_*s*_; 3) the transcribed target gene mRNA M then goes through translation and maturation to form the mature green fluorescent protein (*G*_*m*_), whose intensity is measured to represent the expression level. *α*_*m*_ is the target gene transcription rate, *α*_*gm*_ is the protein maturation rate, *K*_*i*_ is the elongation rate, *K*_*e*_ is the translation rate, and *δ* is the degradation rate of the RNAs.

**Figure 1.**
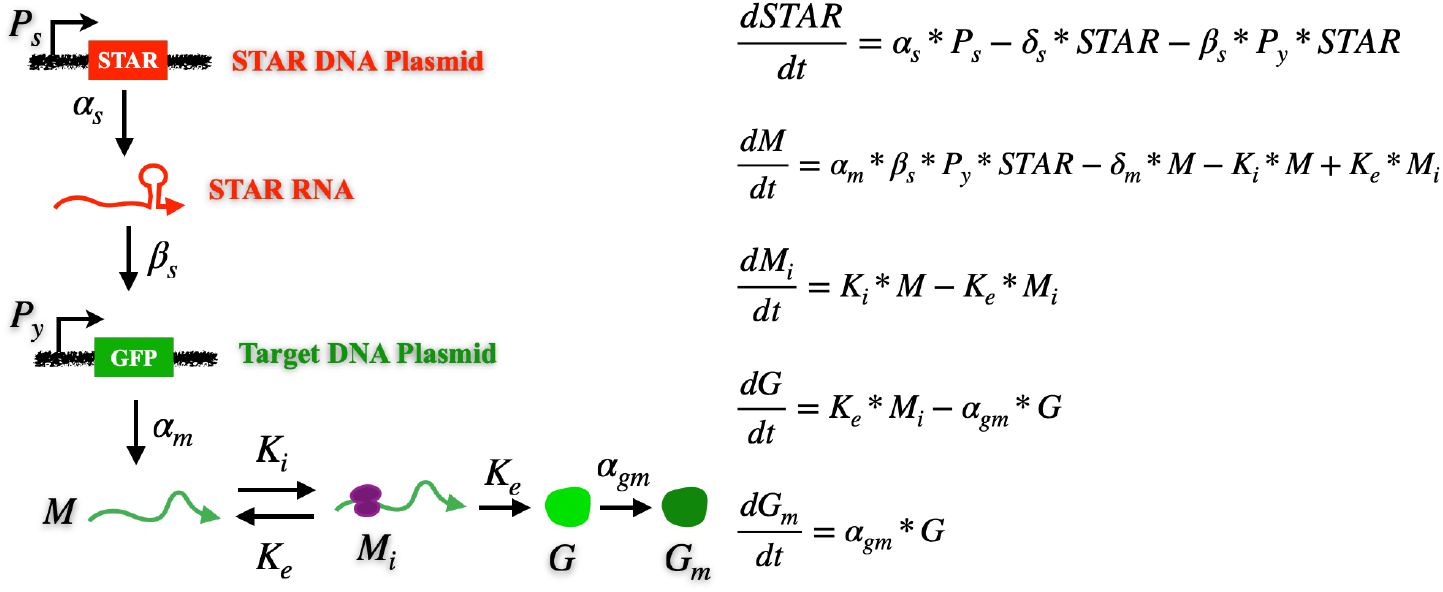
The STAR system activates gene expression by opening the secondary structure that blocks transcription. The simplified process is modeled with and ODE-based mechanistic model to describe the evolution of the key species in the system over time. Figure is modified from Ref. [30] © 2019 Wiley Periodicals, Inc.

Fig. 2a shows the time series enhanced green fluorescent protein (EGFP) concentration measured from the cell-free experiments of the STAR circuit, with different initial STAR DNA Plasmid *P*_*s*_ concentrations. Each condition is independently repeated for nine times for statistical significance, and the EGFP expression level is measured every 5 mins for a total of 240 mins experiment duration. The thicker plots show the average of the nine repeats, whereas the shaded areas show the variation of the measurements. As the *P*_*s*_ concentration increases from 4 nM (blue) to 8 nM (red), the EGFP expression level increases, as more activator is produced in the system. However, due to plasmid *P*_*y*_ saturation, less significant increase from a *P*_*s*_ concentration of 8 nM to 16 nM (yellow) is obtained. Analysis on the EGFP variance across the nine repeats for each *P*_*s*_ concentration indicates a monotonic overall increase over time (Fig. 2b), whereas the ratio of variance/average expression level remains relatively constant over time (Fig. 2c). These observations indicate a time-dependent, nonnegligible measurement variation in the experiments, motivating the incorporation of measurement variance in model parameterization. Note that, all the experimental measurements presented in this study are from Dr. Julius Lucks’ lab at Northwestern, and the details of all the experiments are provided in Ref. [30].

**Figure 2.**
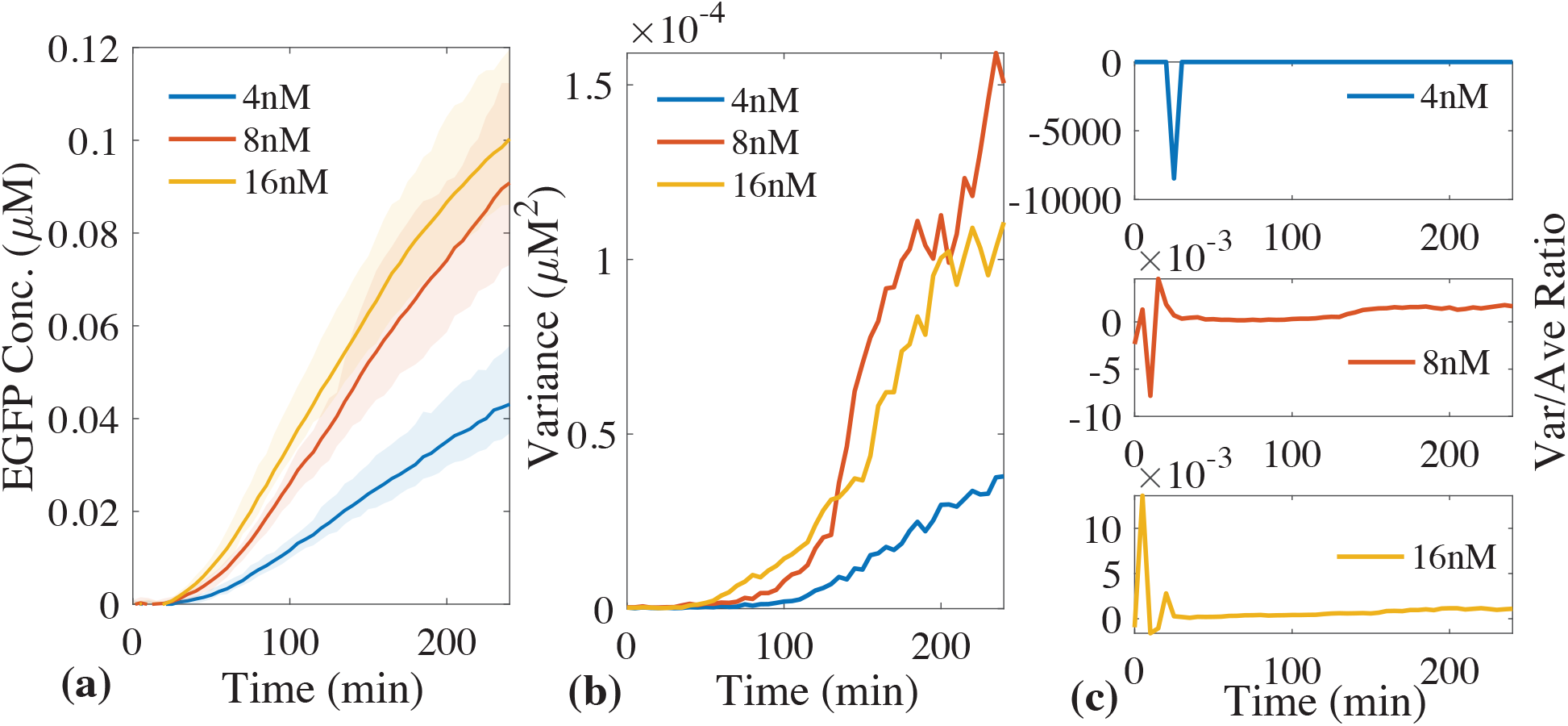
EGFP cell-free experimental measurements for the STAR system with STAR plasmid concentration *P*_*s*_ of 4 nM (blue), 8 nM (red), and 16 nM (yellow). Thick plots in (a) represent the average of nine independent experiment, whereas the shaded area represents the variance across all nine experiments. (b) measurement variance showing a time-dependent, monotonic increase trend for the three conditions. (c) variance over average EGFP level suggests a constant ratio after the initial stage of the experiment.

To investigate the impact of the nonnegligible measurement variance on model parameterization, we first sought to define the “True” kinetic parameters used in the STAR model, as a reference. This set of true parameters is identified by fitting the model to the averaged expression of the *P*_*s*_ = 8 nM condition experiments, using a genetic algorithm, [31–33] with an unweighted pair-wise mean squared error objective function, i.e. 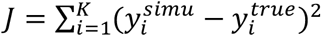, where 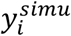 and 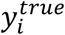 are the simulated and experimentally measured EGFP concentration at discrete time *i. K* = 49 corresponding to the 240 mins of experimental duration. Note that, this set of “True” kinetic parameters should not be confused with the biologically true kinetic parameters, as they would only capture the dynamics of the measured EGFP data and are used to generate a controlled set of synthetic data for our analysis. The specific value of each parameter is given in Supplementary Information SI Table 1.

With the “True” kinetic parameters, we then conducted a simulation for the *P*_*s*_ = 8 nM STAR experiment condition. To introduce variance into the simulation, we first defined the maximum variance value as 20% of the maximum EGFP concentration observed during the 240 mins of the simulation. We then evenly discretized the variance range, i.e. from 0 to the maximum variance value into 48 intervals, with each interval resembling a 5 mins experimental time. We then used a Gaussian distribution with the discretized variance as the standard deviation at each time point, to generate a random variance to the simulated EGFP expression level. Following this mechanism, we generated a total of 100 sets of simulations, with each set containing 3 repeats, mimicking the standard experimental practice. Moving forward, our analysis will focus on these 100 sets of synthetic STAR system data.

## 4. Results

### Analysis with the Synthetic STAR System Dataset

While various optimization algorithms are available for parameterization, from simple gradient-based algorithms [34–37] to advanced probabilistic optimizations, [38–43] here we adopt genetic algorithm, a stochastic optimization algorithm, considering its enhanced capability of identifying the global optimal solution. [33,44–47] Specifically, we used MATLAB command *optimoptions*(‘*ga*’) for all optimization in this study, and we consider three variations of the mean squared error-based objective function: 1) the summation of unweighted mean squared error as in Eqn. 1, named as “unweighted” for short in the following text, and “UnW” in the Figures; 2) the summation of weighted mean squared error, using the inverse variance as in Eqn. 2, named as “v-weighted” for short in the following text, and “Wt” in the Figures; and 3) the summation of weighted mean squared error, using the inverse of the stabilized variance calculated as the following, named as “sv-weighted” for short in the following text, and “WtS” in the Figures.

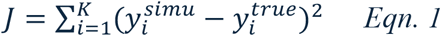

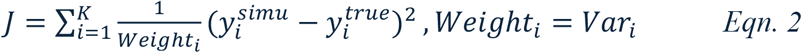

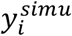 and 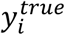 are the simulated and the average (of three repeats) true EGFP concentration (i.e. the synthetic dataset), at each discrete time *i. Var*_*i*_ is the variance of 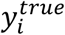 at time *i*. Note that, since the output GFP concentration initiates from 0, to avoid invalid division, *i* starts from time point 1 instead of time 0 in the calculation of the objective function.

The variance of 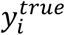 needs to be estimated from the three repeats, and we propose a two-step approach for the variance function estimation. First, we estimate the expected value of *Y*:

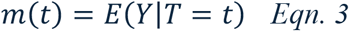

by minimizing *Eqn. 1*. Then, we estimate the variance function as:

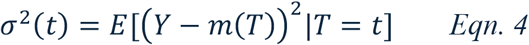

More precisely, let *z*_*tj*_ = (*y*_*tj*_ − m(*t*))^2^ be the squared residuals, the variance function σ^2^(*t*) can then be estimated using nonparametric regression on these squared residuals over time *T*. To obtain a stable estimate, kernel regression or local polynomial regression methods may be applied. [48] Since the data are observed at discrete time points and σ^2^(*t*) is expected to increase over time, we use isotonic regression on the squared residuals *z*_*tj*_, as isotonic regression is a nonparametric technique that estimates a function while preserving a specified monotonic relationship. [49] In our setting, we seek the function σ^2^ that minimizes the squared error loss:

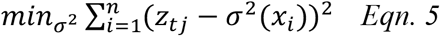

subject to the monotonicity constraint:

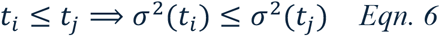

The above-described monotone regression is then achieved using R command *monoreg* from package *fdrtool*. [50] With these specifications, we then parameterized the model using the 100 sets of synthetic repeats. Note that, for all the fittings in this study, each parameter is initiated randomly from within its corresponding range, identified from literature for biological relevance. The specific range of each parameter is summarized in Supplementary Information Table 1 and 2 for the STAR and the CRISPRi system respectively.

Fig. 3 summarizes the fitting results with the three objective functions. The thick plots in Fig. 3a-c represent the average of the 100 simulations for the synthetic dataset (black), predictions with parameters fitted using the unweighted objective function (blue, UnW), and predictions with parameters fitted using the v-weighted objective function (green, Wt), and using the sv-weighted objective function (red, WtS). In terms of the averaged fitting results (thick plots), all three objective functions have led to satisfactory fitting results. However, fitting with the two weighted objective functions yielded a much accurate approximation of the true data at the early stage of the process, with increased variance (shaded area) over time, as compared to that without weighting. This makes sense as the variance in the synthetic dataset is monotonically increasing, and weighting with the inverse variance tends to emphasize more on fitting the early-stage dynamics. Interestingly, weighting with the stabilized variance significantly reduced the variance of the fitting, comparing Fig. 3b and c, providing an improved performance. Further investigation into the variance reveals that, with stabilization the raw variance for adjacent time points now has the same value, thus being weighted the same in the objective function, as the stabilization tends to linearize the variance to reduce noise. Such effect further resulted in shared similarity with the unweighted objective function and presents a mitigation between the unweighted and the raw variance-weighted objective functions. Supplementary Information Fig. 1 (SI Fig. 1) presents the comparison of the raw and the stabilized variance for four randomly selected sets, showing the removal of noise with the stabilization, with a linear approximation.

**Figure 3.**
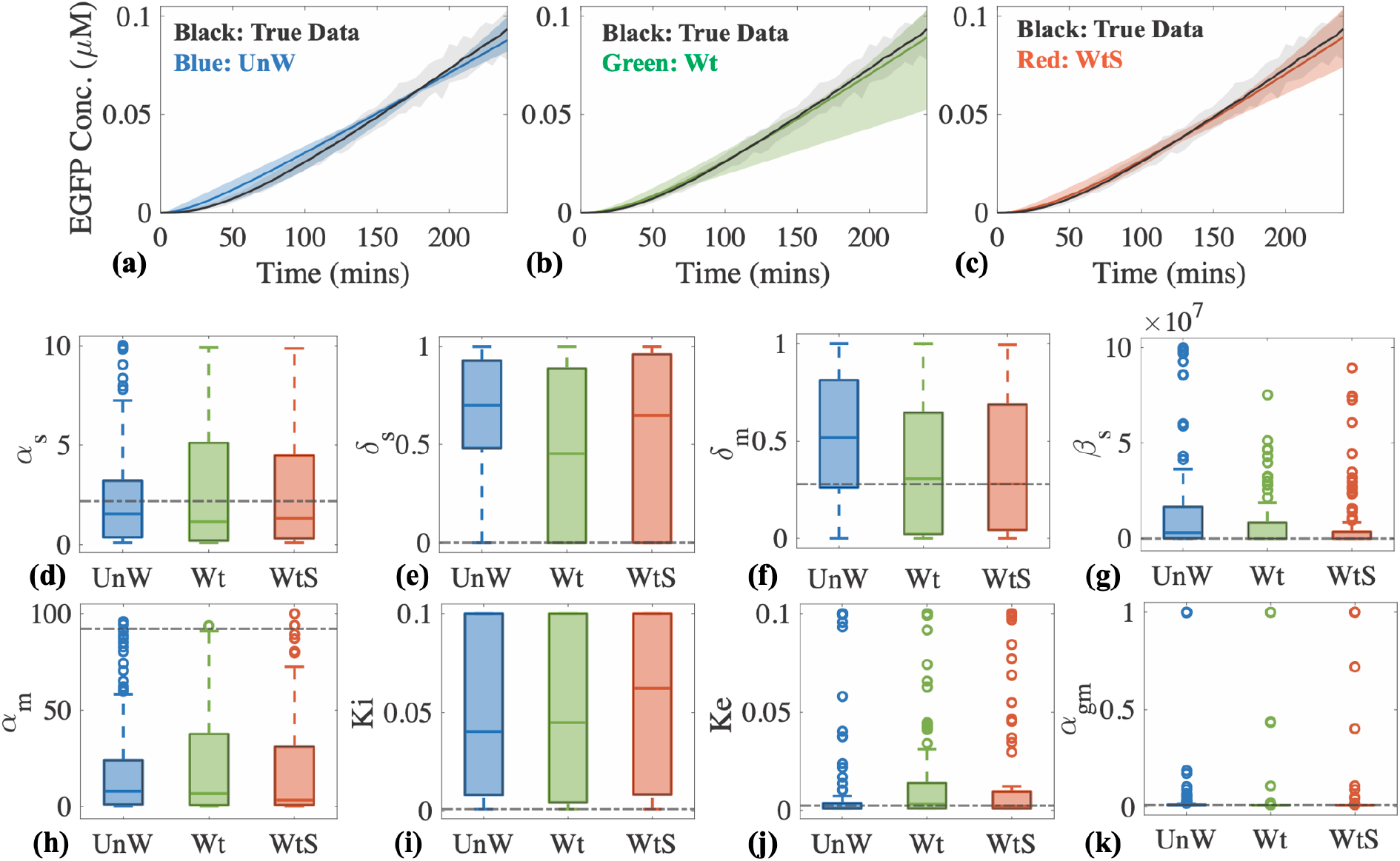
Summary of the fitted results for the synthetic STAR dataset. (a)-(c) comparison of the true EGFP data (black) with the fitted simulations from unweighted (blue, UnW), weighted with variance (green, Wt), and weighted with stabilized variance (red, Wts). (d)-(k) Boxplots showing the distribution of parameters by fitting to the 100 sets of synthetic data. Dashed black horizontal line indicates the true value for the corresponding parameter.

Fig. 3d-k show the distribution of the fitted parameters, with the bar in the box representing the median, the bottom and the top edges of the box representing the 25% and 75% percentiles, and the circle data points representing outliers. The horizontal dashed black line represents the true kinetic value for the corresponding parameter. Comparing the median and the true parameter values, the results reveal that none of the three objective functions successfully identified the correct value for δ_s_ (Fig. 3e), α_m_ (Fig. 3h), and K_*i*_ (Fig. 3i), whereas noticeable improvement was obtained for δ_m_ (Fig. 3f) with the two weighted objective functions. Pair-wise statistical *t*-test in Fig. 4a and b further indicate significant differences in the fitted values for parameters *δ*_*s*_, *δ*_*m*_, and *β*_*s*_, with and without weighting (UnW vs. Wt, and UnW vs. WtS), whereas no significant differences were observed with or without stabilizing the variance (Wt vs. WtS). Note that value 1 in Fig. 4a indicates rejection of the null hypothesis that the two sets of values are from the same distribution with the same mean values, and Fig. 4b presents the *p*-values of the corresponding *t*-test in Fig. 4a.

**Figure 4.**
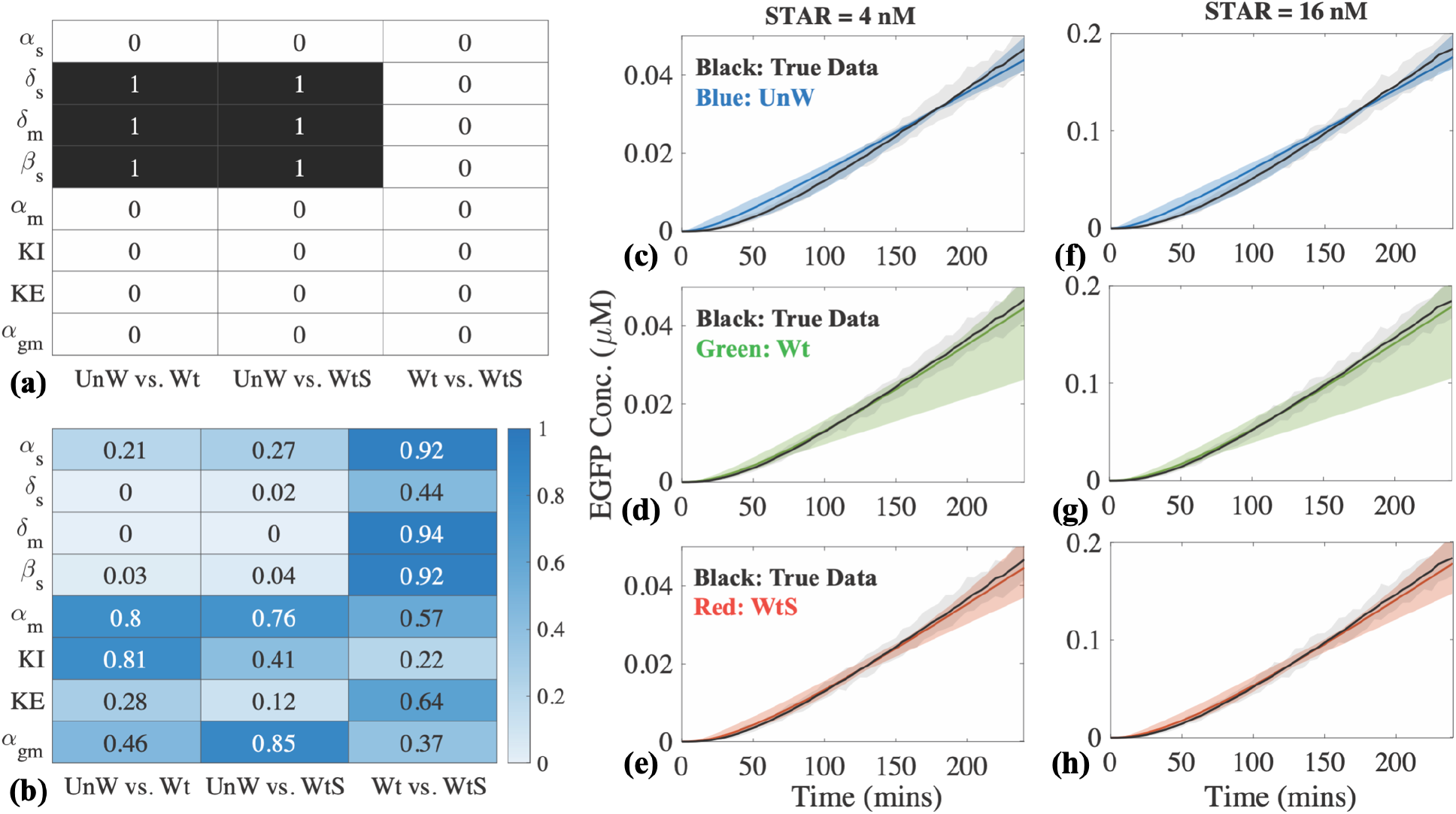
(a)-(b) *t*-test and *p*-values on fitted parameters, showing significant differences in parameters δ_s_, α_m_, and K_*i*_, with and without weighting the objective function. (c)-(h) prediction results for STAR plasmid conditions *P*_*s*_ = 4 nM and *P*_*s*_ = 16 nM, showing that weighted objective functions capture better the early-stage dynamics, and weighting with stabilized variance is able to reduce the variance towards the end of the process. Thick plots represent the average of the 100 individual simulations, and the shaded areas represent the variance across all the simulations.

We then sought to evaluate the accuracy of using the fitted parameters in predicting new conditions, i.e. for the *P*_*s*_ = 4 nM and *P*_*s*_ = 16 nM conditions. Fig. 4c-h present the mean and variances of the simulations with the 100 independent parameter sets fitted with the three objectives in comparison to the true data (i.e. synthetic dataset, in black). Similar observations as in the fitting results in Fig. 3 can be made that, parameters fitted from the weighted objective functions (green, Wt, and red, WtS) provided a much higher accuracy for early-stage dynamics, with an increased variance towards the end of the process, as compared to that without weighting (blue, UnW). Again, weighting with the stabilized variance mitigated the undesirably large variance towards the end.

Based off these findings, we hypothesize that, for a system with measurements showing a monotonically or an overall increasing variance, weighting the pair-wise squared error by the inverse stabilized measurement variance could provide satisfactory estimates of the model parameters. We then set to evaluate this hypothesis by focusing on a CRISPRi system, which contains more complex interactions than the STAR system.

### Hypothesis Evaluation with the CRISPRi System Experiments

The same CRISPRi system as in Ref. [30] is considered here, and summarized in Fig. 5a. *crRNA* and *tracrRNA* mRNAs are independently transcribed from plasmids *P*_*cr*_ and *P*_*tr*_ constitutively at rates *α*_*cr*_ and *α*_*tr*_, and can bind to form the guide RNA (gRNA) at rate *γ*_1_, which then binds to *dCas9* to form the repressor complex at rate *γ*_2_. The *CRISPRi complex* then binds to the target gene expression plasmid *P*_*y*_ at rate ω to suppress the translation of EGFP. Fig. 5b shows the time series EGFP concentration measured from cell-free experiments, for initial CRISPRi DNA Plasmids concentrations *P*_*cr*_ = P_*tr*_ = 0.1 nM (blue), 0.25 nM (red), and 0.5 nM (yellow). Each condition independently repeated nine times for statistical significance. The thicker plots are the average of the nine repeats and the shaded areas show the variation of the measurements. Each experiment lasted for 240 mins, with the EGFP measurements taken every 5 mins. Fig. 5c shows the variance and Fig. 5c shows the variance over averaged expression level for each condition over time. Notice that the measurement variances for all three conditions showed an overall increasing trend (Fig. 5c), with the 0.25 nM experiments showing a pronounced spike at the early stage, rather than a strict monotonic increase as in our synthetic STAR dataset.

**Figure 5.**
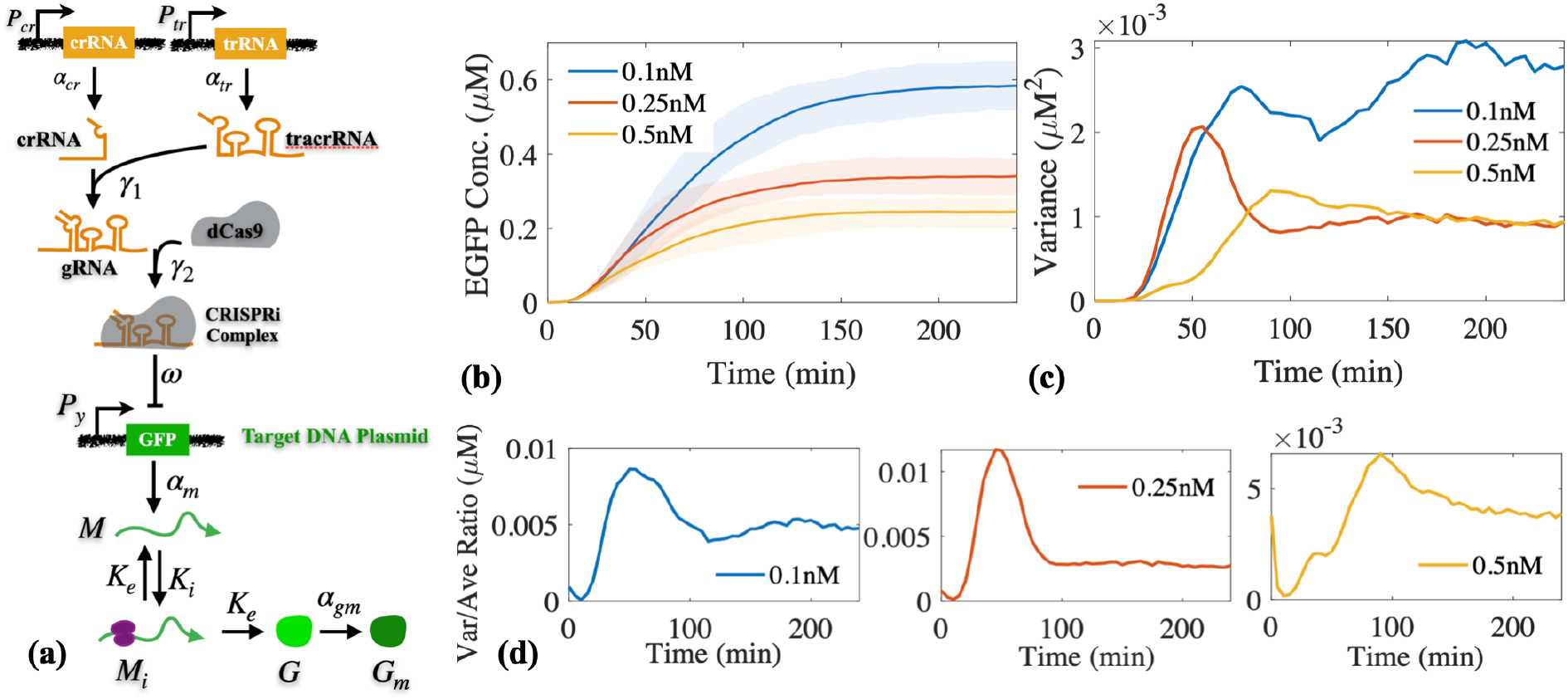
(a) Diagram showing the key reactions in the CRISPRi system, with associated kinetic parameters. (b) Cell-free experimental EGFP expression measurement for *P*_*cr*_ = P_*tr*_ = 0.1 nM (blue), 0.25 nM (red), and 0.5 nM (yellow). (c) Measurement variances show an overall increase trend in all three conditions, except for a prominent pulse profile for the 0.25 nM condition. (d) Variance/average ratio time trajectories show a relatively constant relationship between the variance and the average expression level over time. Figure (a) is modified from Ref. [30] © 2019 Wiley Periodicals, Inc.

The same ODE model from Ref. [30] is again used for our analysis and is presented in Fig. 6. We first fitted the model to the average EGFP expression level from the 0.25 nM-conditioned experiments, to get kinetic parameter values to predict the 0.1 nM and 0.5 nM conditions. Fitting results in Fig. 6a indicate all three objective functions yielded satisfactory approximation to the averaged experimental measurement (thick black plot). Surprisingly, predictions using the fitted parameters suggest, that parameters fitted with the v-weighted objective function yielded the worst prediction accuracy (green plot in Fig. 6b and c), whereas parameters fitted with both unweighted (blue) and vs-weighted (red) objective functions yielded satisfactory accuracy, with the vs-weighted providing a slightly better performance. These findings suggest that while weighted objective function could improve the model parameterization, the specific design of the weighting can dramatically affect the performance, and weighting with the stabilized measurement variance could be a reliable choice for gene circuit model parameterization.

**Figure 6.**
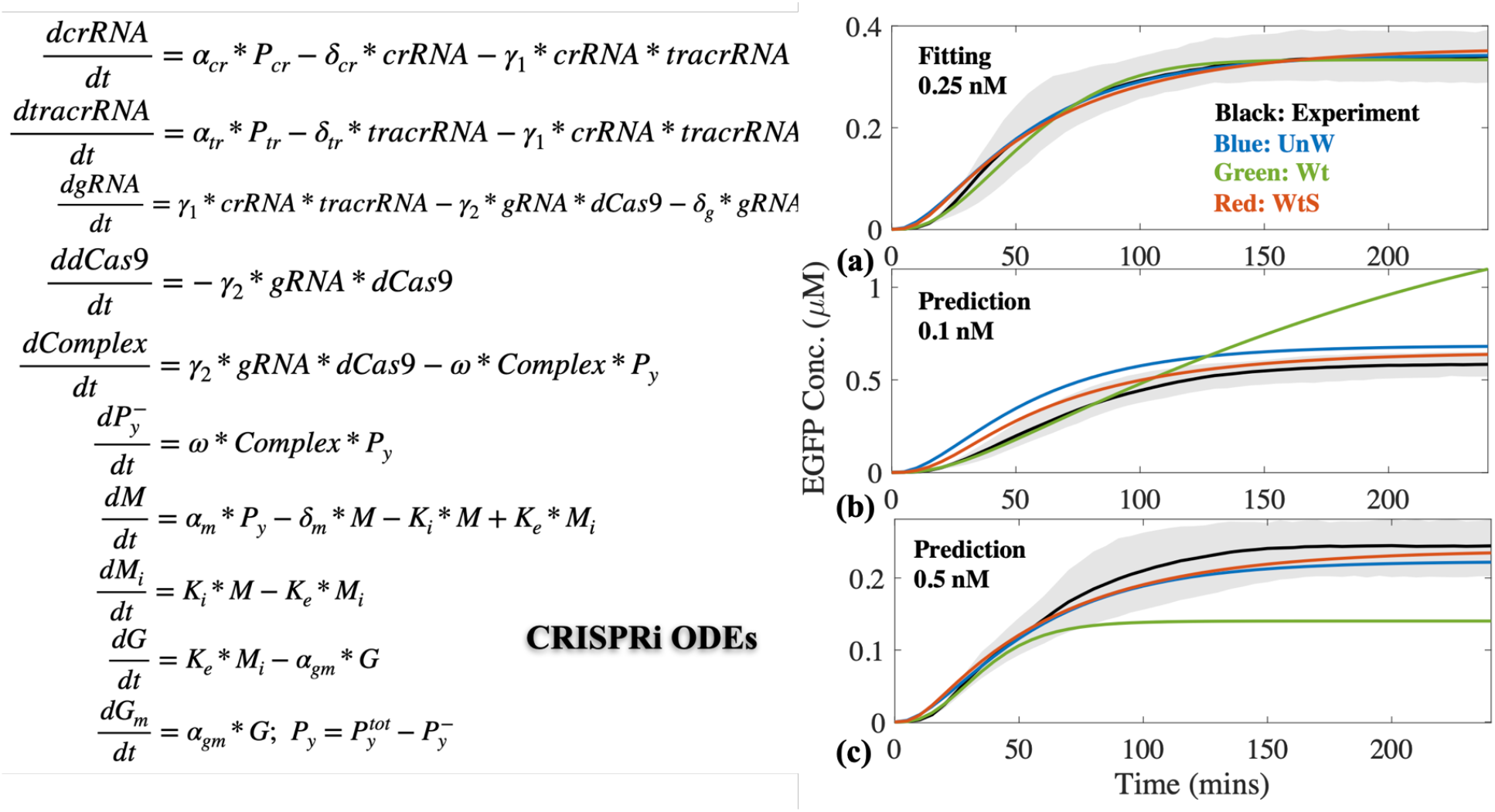
ODE model of the CRISPRi system. (a) Fitting results to CRISPRi 0.25 nM condition experiments. Thick black plot represents the average of the nine independent experiments, with shaded area representing the variance. Blue plot represents fitting with the unweighted objective function, green plot for fitting with inverse variance weighted objective function, and red plot for fitting with inverse stabilized variance weighted objective function. Same color scheme applies to (b) and (c). (b) Predictions with parameters fitted from the 0.25 nM experiments for the 0.1 nM condition. (c) Predictions with parameters fitted from the 0.25 nM experiments for the 0.5 nM condition.

## 5. Conclusion and Discussion

Mathematical modeling has become an integral tool in systems and synthetic biology to probe the fundamentals of biological systems, to thus design remedies to diseases. The accuracy of a mathematical model can be affected by its underlying assumptions, complexity, structure, and associated model parameters. In this study, we investigated how the design of the objective function, in terms of incorporation of the measurement variance, would affect the accuracy of ordinary differential equation-based model parameterization. By analyzing a STAR and a CRISPRi synthetic gene circuit, with cell-free experimental measurements, we found that weighting the pair-wise squared error with the stabilized measurement variance could provide more accurate approximation of the dynamics, with mitigated prediction variance, as compared to that without weighting or weighted with the non-stabilized variance, if the measurements present a monotonic or generally increasing trend over time. Surprisingly, fitting with the unweighted objective function in general has yielded satisfactory results, despite slightly inferior accuracy for the early stage approximate and capturing the variance of the system measurement. However, given the fact that all models are built upon assumptions

While promising, we also noticed several issues during the study. First, the fitting analysis with the synthetic STAR dataset indicates that some of the true kinetic parameters were not correctly identified with any of the three tested objective functions. This could be because there exist multiple solutions to the ODEs to approximate the output expression level, that more and specifically designed experiments are necessary to identify a specific parameter. It could also be a result of the oversimplification of the reactions in the model, that multiple reaction rates are lumped together into one parameter to capture the overall dynamics. Second, to our surprise, when weighted with the stabilized variance, we noticed significant increase in the computational time with the genetic algorithm used here, from the order of minutes to the order of hours. However, this phenomenon is not observed in the CRISPRi system. In-depth investigation on this issue is needed, to understand if this is due to the genetic algorithm or the model construction or the dataset in our study. Third, while genetic algorithm provides an enhanced possibility of finding the global optimum of the solution, we noticed that it could take a significant amount of time and iterations for one optimization to converge to a solution, whereas seconds or mins for some other trials. Such as huge variation in computational cost could construct a drawback for system with multiple local optima, therefore the trustiness of the fitted parameters, that alternative and more robust optimization approach should be considered. Fourth, while measurement variance is unavoidable and prevalent in biological experiments, the pattern of experimental variance could vary from system to system and could also be hard to identify. To establish a generally applicable guideline for incorporating measurement variable in model parameterization would require extensive validation for a wide range of circuits and systems, with increased complexity.

Given the increasing importance of mathematical modeling in systems and synthetic biology, we anticipate continued efforts on the understanding of measurement variance on efficient and accurate model parameterization.

## Ethical Statement

No applicable.

## Data Accessibility

All the MATLAB scripts used for this study are uploaded to GitHub, accessible at: https://github.com/xtang38/Thales-et-al-Objective-Function-2025.git

## Authors’ Contributions

T.R.S.: acquisition of data, analysis and interpretation of data, drafting and revising the article.

W.T.: conception and design, acquisition of data, analysis and interpretation of data, drafting and revising the article.

X.T.: conception and design, acquisition of data, analysis and interpretation of data, drafting and revising the article.

