## Supplementary Document for "Investigating the Impact of Measurement Variance on Gene Circuit Model Parameterization"

**SI Table 1.** Kinetic parameter information for STAR system synthetic data generation, true values are from STAR experiment with $P_s = 8$ nM condition, parameters in all fittings are confined within the specified ranges.

| Parameter | Description | True Value | Range for Fitting |
|---|---|---|---|
| $\alpha_s$ | STAR RNA transcription rate | 2.198 $s^{-1}$ | 10e$^{-2}$ ~ 10 $s^{-1}$ |
| $\delta_s$ | STAR RNA degradation rate | 3.484e$^{-4}$ $s^{-1}$ | 10e$^{-5}$ ~ 10e$^{-1}$ $s^{-1}$ |
| $\delta_m$ | EGFP mRNA degradation rate | 0.279 $s^{-1}$ | 10e$^{-5}$ ~ 10e$^{1}$ $s^{-1}$ |
| $\beta_s$ | STAR binding rate | 1.004e$^{3}$ $nM^{-1}$ | 10e$^{3}$ ~ 10e$^{7}$ $nM^{-1}$ |
| $\alpha_m$ | Lumped STAR activation and transcription rate | 92.231 $s^{-1}$ | 10e$^{-2}$ ~ 10e$^{2}$ $s^{-1}$ |
| $K_i$ | EGFP translation initiation rate | 0.001 $s^{-1}$ | 10e$^{-4}$ ~ 10e$^{-2}$ $s^{-1}$ |
| $K_e$ | EGFP translation elongation rate | 0.0025 $s^{-1}$ | 10e$^{-4}$ ~ 10e$^{-2}$ $s^{-1}$ |
| $\alpha_{gm}$ | EGFP maturation rate | 0.011 $s^{-1}$ | 10e$^{-3}$ ~ 10e$^{-1}$ $s^{-1}$ |

**SI Table 2.** Kinetic parameter information for CRISPRi system

| Parameter | Description | Range for Fitting |
|---|---|---|
| $\alpha_{cr}, \alpha_{tr}$ | CRISPR/Tracr RNA transcription rate | $10e^{-2} \sim 10\ s^{-1}$ |
| $\delta_{cr}, \delta_{tr}$ | CRISPR/Tracr RNA degradation rate | $10e^{-5} \sim 10e^{-1}\ s^{-1}$ |
| $\delta_m$ | EGFP mRNA degradation rate | $10e^{-5} \sim 10e^{1}\ s^{-1}$ |
| $\gamma_1$ | crRNA/trRNA binding rate | $10e^{3} \sim 10e^{7}$ $nM^{-1}$ |
| $\gamma_2$ | gRNA/dCas9 binding rate | $10e^{3} \sim 10e^{7}$ $nM^{-1}$ |
| $\omega$ | CRISPRi/plasmid binding rate | $10e^{3} \sim 10e^{7}$ $nM^{-1}$ |
| $\alpha_m$ | Lumped STAR activation and transcription rate | $10e^{-2} \sim 10e^{2}\ s^{-1}$ |
| $K_i$ | EGFP translation initiation rate | $10e^{-4} \sim 10e^{-2}\ s^{-1}$ |
| $K_e$ | EGFP translation elongation rate | $10e^{-4} \sim 10e^{-2}\ s^{-1}$ |
| $\alpha_{gm}$ | EGFP maturation rate | $10e^{-3} \sim 10e^{-1}\ s^{-1}$ |

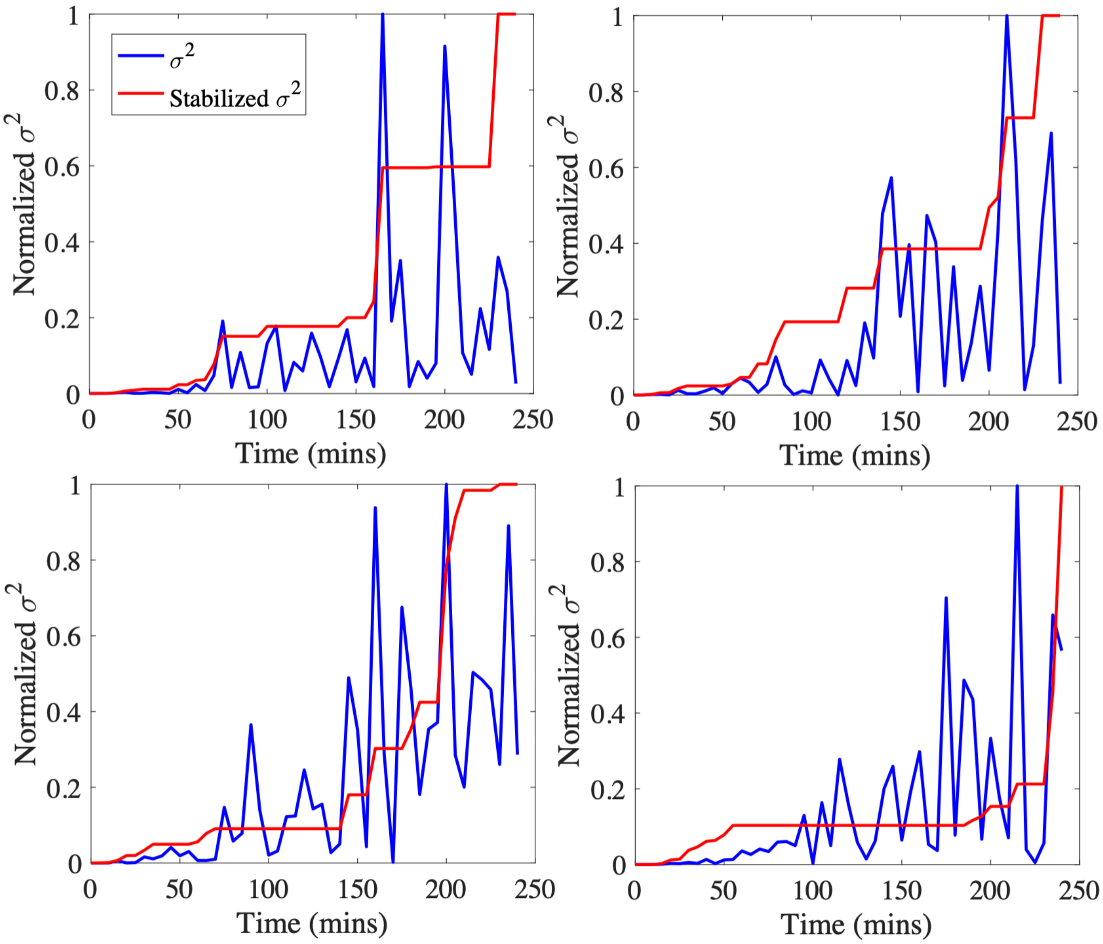


**SI Figure 1.** Comparison of four randomly selected raw variance and stabilized variance time trajectories from the synthetic STAR system dataset, showing the stabilization tends to linearize the raw variance to remove noise.
